# Fully automated open-source analysis and interactive visualization of magnetic resonance spectroscopic imaging (MRSI) data in Osprey-MRSI

**DOI:** 10.64898/2026.09.02.748934

**Authors:** Helge J. Zöllner, Alexander R. Craven, Vasilis M. Karlaftis, William T. Clarke, Dillip K. Senapati, Doris D.M. Lin, Georg Oeltzschner, Peter B. Barker

## Abstract

**Purpose:** Magnetic resonance spectroscopic imaging (MRSI) is a versatile technique to investigate the spatial distribution of *in vivo* metabolism. However, processing MRSI data is demanding, and only a few software packages support end-to-end analysis. The goal of this study was to implement fully automated, end-to-end MRSI analysis into the open-source ‘Osprey-MRSI’ software package.

**Methods:** MRSI-specific analysis and visualization capabilities were implemented, building on the existing Osprey workflow. Modifications included spatial transformation and filtering operations, automated brain masking and tissue segmentation of the MRSI data, improved lipid filtering, rapid integral maps, linear-combination modeling with explicit B_0_ frequency-shift correction, and generation of quality-control maps and metabolic images. A fully interactive GUI and semi-interactive HTML reports provide a user-friendly way to inspect each step of the analysis. All analysis derivatives are also exported in NIfTI and NIfTI-MRS format for easy visualization and synergies with other toolboxes and modalities.

**Results:** The automated MRSI workflow was successfully used to analyze short- and medium-TE 3T *in vivo* MRSI datasets from all major vendors (Philips, GE, Siemens) across multiple sites. Correct coregistration of MRSI data and MR images was validated using phantom data from each vendor and existing MRSI processing tools.

**Conclusion:** Osprey-MRSI offers state-of-the-art methods with minimal user interaction available for non-expert users. The modularity of the workflow and the modeling algorithm will foster innovation and development of novel MRSI-specific analysis methods.

## Introduction

Magnetic resonance spectroscopic imaging (MRSI) is one of the few methods for mapping of metabolite levels in the human brain ^1^. However, MRSI analysis is more demanding than single voxel (SV) magnetic resonance spectroscopy (MRS) because it involves processing many spectra and more variable spectral quality. Only a few existing software packages support end-to-end analysis^2–7^. Often, the underlying processing code is simply a modified version of an SV processing pipeline, even though MRSI poses its own set of challenges. Modified SV methods typically do not consider spatial information, which limits one of the advantages of MRSI, for example, when prior spatial information is included during the removal of nuisance lipid signals. The large number of MRSI voxels (>7500 for high-resolution MRSI with extended spatial coverage) poses a similar computational challenge, which can be addressed through hardware parallelization, a feature often overlooked in SV analysis and therefore rarely implemented. Similarly, visualization of MRSI data is a challenge best addressed by an interactive display that simultaneously displays the MR spectrum and its origin (i.e. spatial location) on an MR image. Finally, for analyzing multiple brain regions, the most powerful approach is atlas-based statistics, as it includes most MRSI voxels, reduces bias, and minimizes the risk of missing changes across the brain. While some of the features described in this paragraph are implemented in currently available software packages^2–7^ or in in-house scripts^8^ used by research groups focused on MRSI method development, broad access for non-expert users to these packages remains limited, thereby hampering the application of MRSI in research and clinical settings despite its potential. **Supplementary Material 1** presents currently available MRSI analysis^2–7,9–15^ software, along with their features, programming language, and code availability.

A brief overview of selected software tools and their features is provided below. The Metabolite Imaging and Data Analysis System (MIDAS) was designed to analyze MRSI data collected using the echo-planar spectroscopic imaging (EPSI) sequence ^2^. MIDAS includes MRSI-specific preprocessing, such as brain and lipid masking. It performs spectral analysis using a linear-combination model. Finally, it uses automated quality evaluation, tissue-fraction correction, spatial normalization to create a study database, and atlas analysis to visualize high-dimensional MRSI data in a digestible way. MIDAS is the most developed MRSI workflow to date; however, the long-term sustainability of the MIDAS package, including its use of the IDL language and its adaptation for non-EPSI applications, remains uncertain. FSL-MRS is a Python-based software package that provides consensus-recommended^16^ end-to-end analysis of MRS and MRSI data^6^, which uses the NIfTI-MRS community standard^17^ for data import and export. Lipid masking and segmentation can be performed using FSL’s algorithms. Spectral analysis is performed with its flexible linear-combination algorithm^18^. It interfaces with the NIfTI-Viewer FSLeyes via a plugin, enabling fully interactive inspection and visualization of the MRSI data^17,19^. LCModel^20^, the *de facto* gold-standard linear-combination algorithm in the single-voxel MRS, is also able to process MRSI data. It allows Bayesian priors for first-order phase corrections and frequency shifts and analyzes the data from the center voxel outward. The model outputs are visualized in a static file, whereas metabolite maps must be generated with a separate script, making output visualization challenging.

Osprey is a MATLAB-based ^1^H-MRS analysis package that currently offers end-to-end analysis of SV data for all major vendors, with increasing adoption by the MRS community ^21^. Building on the FID-A package _22_, it has been consistently maintained and expanded, with new features including a fully automated, compiled workflow for servers and Docker containers and a flexible, generalized linear-combination model that supports new model functions and multidimensional modeling.

The primary objective of this study is to present the Osprey-MRSI workflow, including preprocessing, linear-combination modeling, quantification, and atlas-based statistics. Second, we demonstrate the interactive workflow features and interfacing with other neuroimaging toolboxes using several MRSI datasets.

## Methods

### Building the Osprey-MRSI Analysis Workflow

Osprey is a MATLAB-based, open-source MRS software package that provides end-to-end analysis of single-voxel MRS data, following expert consensus recommendations^21^. We modified each of Osprey’s eight native modules to support MRSI data for automated analysis of all MRSI voxels (see **Table 1**), building on top of Osprey version 3.0.0. The workflow is available on GitHub (https://github.com/HJZollner/osprey-mrsi), additional data and descriptions are available on the Open Science Framework^26^. Details are described below, and the Osprey-MRSI analysis workflow is summarized in **Figure 1**.

**Figure 1.**
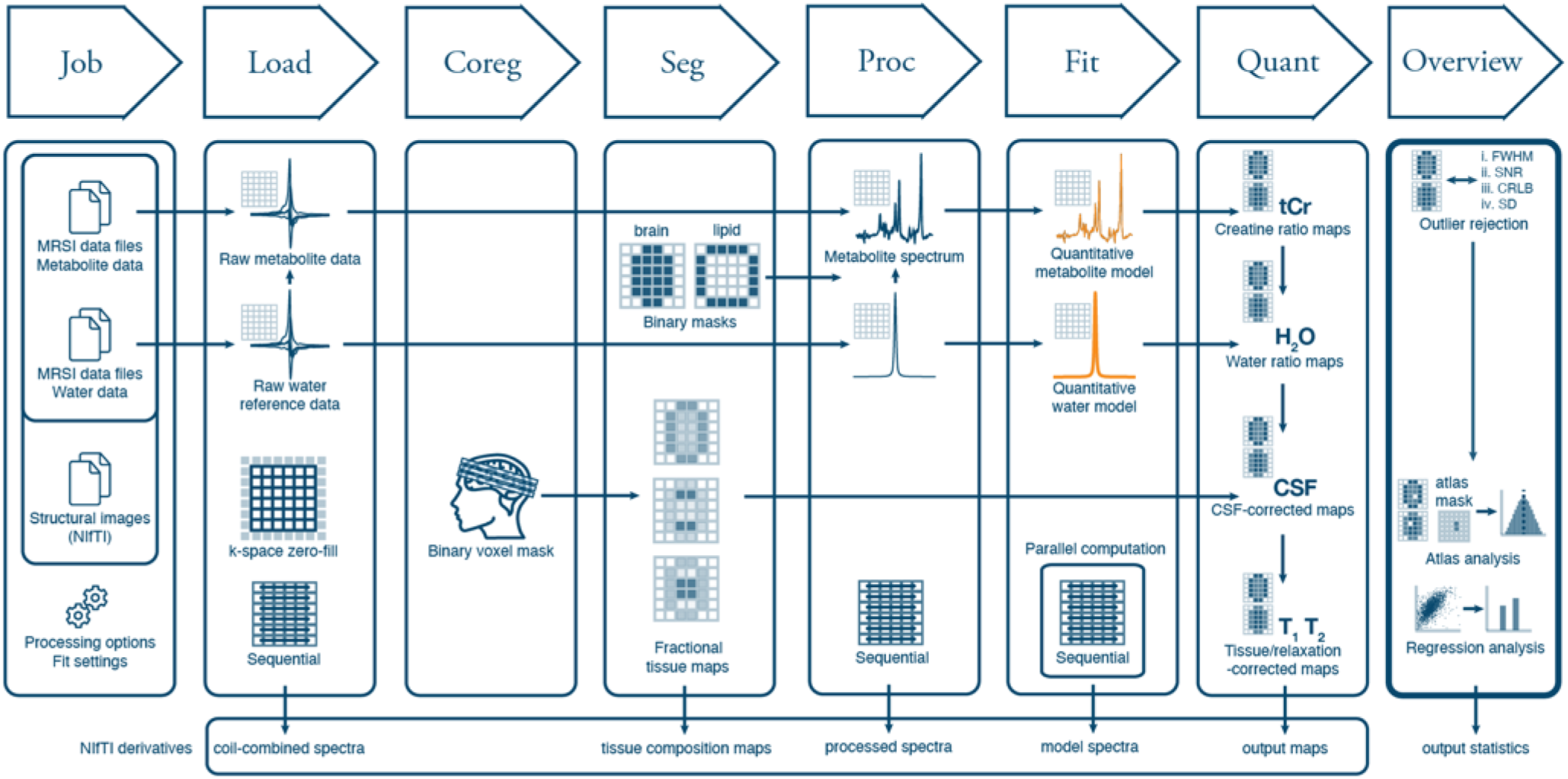
Summary of Osprey MRSI Analysis Workflow. The workflow has eight modules Job, Load, Coreg(istration), Seg(mentation), Proc(ess), Fit, Quanti(fy), and Overview.

**Table 1.**
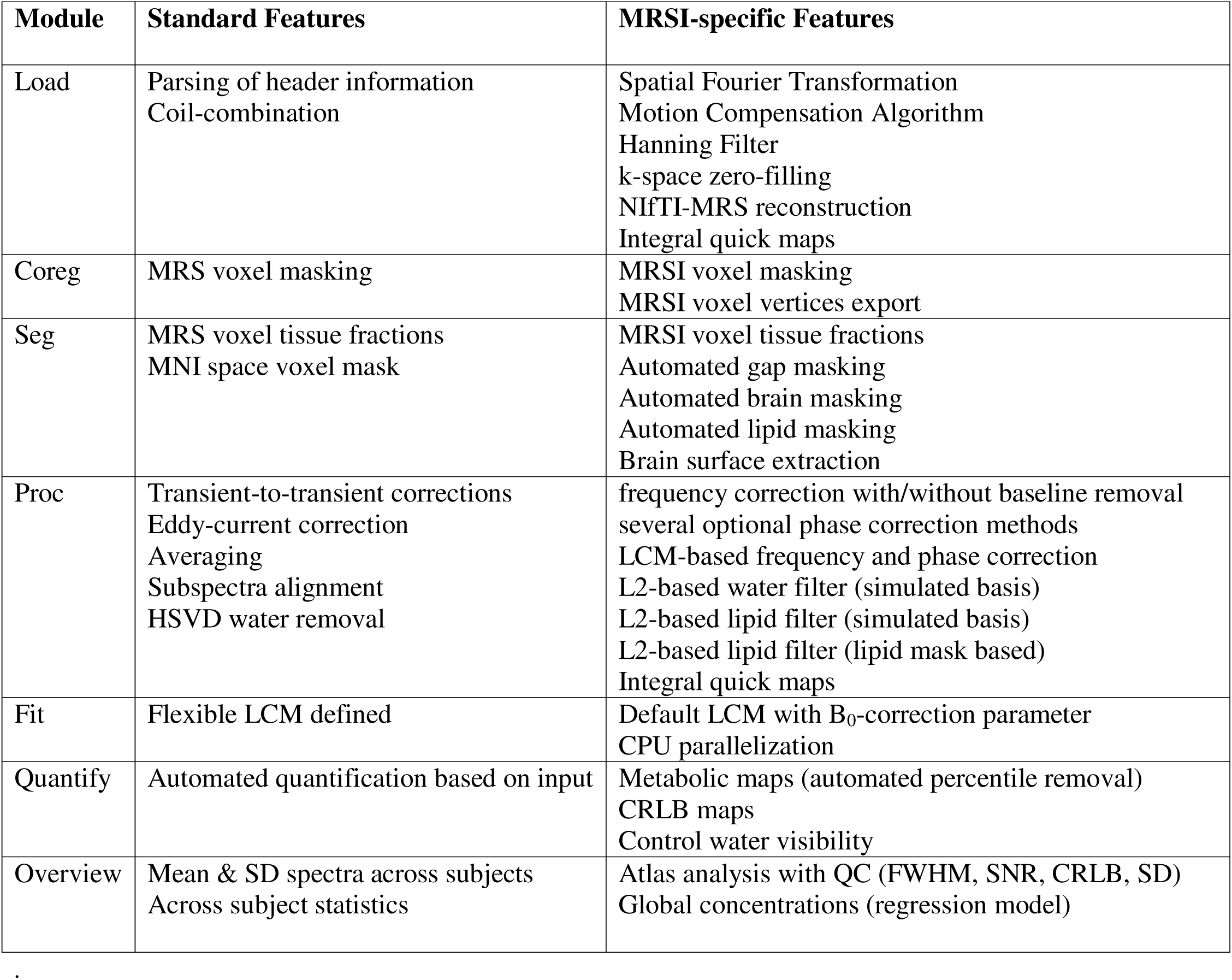
Summary of standard and MRSI-specific features of the Osprey Analysis Workflow.

| **Module** | **Standard Features** | **MRSI-specific Features** |
| --- | --- | --- |
| Load | Parsing of header information<br>Coil-combination | Spatial Fourier Transformation<br>Motion Compensation Algorithm<br>Hanning Filter<br>k-space zero-filling<br>NIfTI-MRS reconstruction<br>Integral quick maps |
| Coreg | MRS voxel masking | MRSI voxel masking<br>MRSI voxel vertices export |
| Seg | MRS voxel tissue fractions<br>MNI space voxel mask | MRSI voxel tissue fractions<br>Automated gap masking<br>Automated brain masking<br>Automated lipid masking<br>Brain surface extraction |
| Proc | Transient-to-transient corrections<br>Eddy-current correction<br>Averaging<br>Subspectra alignment<br>HSVD water removal | frequency correction with/without baseline removal<br>several optional phase correction methods<br>LCM-based frequency and phase correction<br>L2-based water filter (simulated basis)<br>L2-based lipid filter (simulated basis)<br>L2-based lipid filter (lipid mask based)<br>Integral quick maps |
| Fit | Flexible LCM defined | Default LCM with $B_0$-correction parameter<br>CPU parallelization |
| Quantify | Automated quantification based on input | Metabolic maps (automated percentile removal)<br>CRLB maps<br>Control water visibility |
| Overview | Mean & SD spectra across subjects<br>Across subject statistics | Atlas analysis with QC (FWHM, SNR, CRLB, SD)<br>Global concentrations (regression model) |

#### OspreyJob

*OspreyJob* is the interface between the user and Osprey-MRSI. It parses the analysis options, file, and derivative locations from a single input job file. For single-voxel MRS, Osprey supports batch analysis of multiple subjects, e.g., the entire study cohort. Given the increased storage needs of MRSI data, it is recommended to define one job per subject. All relevant workflow derivatives are exported as NIfTI-MRS or NIfTI files.

#### OspreyLoad

*OspreyLoad* loads MRSI raw data, currently with vendor-native support for Philips (.data/.list;.spar/.sdat) and the new community data standard, NIfTI-MRS^17^. NIfTI-MRS files can be generated using the spec2nii tool^17^, which supports virtually every vendor, enabling multi-vendor support for Osprey-MRSI without additional implementation. Note that MRSI data from non-cartesian acquisition types must be fully reconstructed and converted to a NIfTI-MRS file before being passed to Osprey-MRSI. In addition to standard features for parsing header information and performing coil combination, we added several MRSI-specific features. These include optional features such as motion compensation for multi-average data^27^, k-space zero-filling to increase apparent resolution or match resolutions between MRSI scans, Hanning filtering, and spatial Fourier transformation. Integral maps of freely definable frequency regions can be generated, e.g., to rapidly produce metabolic, water, lipid, or water-suppression maps, for early quality assurance.

#### OspreyCoreg

*OspreyCoreg* performs coregistration of the MRSI scan with anatomical images specified by the user. The MRSI-specific features include an MRSI voxel index mask with unique indices for each MRSI voxel, a GIfTI file that defines the MRSI scan as a grid, and, if a multi-slice 2D MRSI was performed, a mask that defines the between-slice gaps. If the user requests interactive processing, which can be useful for *in vitro* data processing, a figure for interactive data masking is shown (see *in vitro* workflow demonstration).

#### OspreySeg

*OspreySeg* performs tissue segmentation based on anatomical MR images. For segmentation, SPM12 generates probability maps for gray matter, white matter, and cerebrospinal fluid, as well as deformation fields between subject and standard space^28^. In addition, a soft matter probability map is generated to identify lipid fractions. The tissue fraction maps at MRSI resolution are computed from the high-resolution probability maps, including accounting for slice gaps in multi-slice 2D-MRSI data. Automated brain and lipid masks are created using user-defined thresholds, and a brain-surface GIfTI file is generated for visualization.

#### OspreyProc

*OspreyProc* performs consensus-guided preprocessing of MRSI data sequentially for each spectrum. The standard features include algorithms for transient-to-transient correction, averaging, eddy-current correction, alignment of subspectra for edited spectroscopy experiments, and optional water removal via Hankel Singular Value Decomposition (HSVD)^29–33^. New MRSI-specific features include options for frequency correction via crosscorrelation, with or without baseline removal; several optional phase correction methods; or LCM-based frequency and phase correction^34^; L2-based filtering of nuisance signals using a simulated basis^35^; or the previously derived lipid mask^36^. L2-based filtering is performed via a single matrix operation across the entire MRSI dataset, making it considerably faster than separate HSVD filtering of each spectrum. Finally, integral maps and quality assurance maps (2-ppm NAA SNR and linewidth) are constructed from user inputs and the processed spectra.

#### OspreyFit

*OspreyFit* performs consensus-recommended linear-combination modeling of the MRSI data^16^ using CPU parallelization to accelerate the procedure. Additionally, the automated (and/or interactive) brain mask reduces the number of MRSI voxels to be fitted. The generalized LCM algorithm used for modeling was recently introduced^25^ and enables a flexible definition of *model procedures*, facilitating adaptation and exploration of new modeling approaches. For example, the default model for MRSI data includes a global frequency-shift term for B_0_-correction^34^. Osprey-MRSI includes model procedures optimized for short-, medium-, and long-TE MRSI data, as well as GABA-edited MRSI data. If supplied, water MRSI data is also modeled using LCM.

#### OspreyQuantify

*OspreyQuantify* computes metabolic maps for all estimated metabolites. The standard features include voxel-based quantification approaches based on user-provided data: raw amplitudes, creatine-referenced estimates, water-scaled estimates, CSF-corrected water-scaled estimates, and fully tissue- and relaxation-corrected molal concentration estimates. Briefly, water-scaling is performed analogously to LCModel^37^. However, the user can define the relative water visibility used during quantification, e.g., for *in vitro* data. CSF-corrected water-scaled estimates correct for the CSF tissue fraction of each MRSI voxel. Fully tissue- and relaxation time-corrected molal concentrations are calculated for each MRSI voxel^38^ from the tissue fraction maps and applies tissue-specific relaxation corrections to each metabolite and water, using literature values^39–43^ for the relaxation times and water visibility from normal adult studies. The code can be modified to account for differences in relaxation or water visibility due to pathology or specific study population.

#### OspreyOverview

*OspreyOverview* performs fully automated statistical and secondary analysis on the final quantitative MRSI outputs. It performs descriptive statistical analysis using standardized brain atlases. Currently, Osprey-MRSI has two atlases available: (1) An atlas combining the AAL3^44^ (138 anatomical labels) and a white-matter atlas^45^ (48 anatomical labels) to a total of 186 anatomical labels separated by brain hemisphere. (2) The neuromorphometrics atlas (Neuromorphometrics, Inc)^46^ with 136 cortical regions and a global white matter mask separated by brain hemisphere. Osprey-MRSI’s atlas workflow produces outputs both separated by hemisphere and combined. The user may provide their own anatomical atlas if it is in MNI standard space. The atlas workflow has four steps: First, the atlas is transformed into subject space. Second, a user-defined fractional threshold generates binary masks for each anatomical label in MRSI space. Third, quality control thresholding of each metabolite and anatomical region is performed using user-defined thresholds, beginning with FWHM and SNR filtering. Then, per-metabolite CRLB filtering can be applied, and finally, per-metabolite outlier rejection is applied, which excludes MRSI voxels that are outside a predefined range, e.g., 3 standard deviations, of the mean. Fourth, the final statistical analysis is performed across all metabolites, quantifications, and anatomical regions, and the results are exported in tabular form.

The *OspreyOverview* module also calculates tissue-specific concentrations using global linear regression, which has been shown to improve assessment of diffuse metabolic changes^47–49^ compared to atlas-based approaches. This approach accounts for partial-volume effects in metabolite estimates and improves effective SNR. The signal of each MRSI voxel is assumed to be a linear combination of the tissue composition of each MRSI voxel and a global single concentration for each tissue type. The tissue fraction maps, along with a user-defined threshold for the sum of gray matter and white matter fractions and slice indices, are used to generate a tissue mask. Next, standard-deviation-based outlier rejection is applied to the metabolite estimates within the mask. Finally, a linear regression analysis estimates tissue-specific concentrations.

### Visualization of MRSI workflow derivatives

While static visualization can assess single-voxel MRS data, interactive visualization is essential to an MRSI workflow.

#### Visualization in Osprey

Osprey-MRSI workflow offers optional interactive visualizations. The workflow can be paused after *OspreyCoreg* and then after each subsequent module. After *OspreyCoreg*, the MRSI grid is superimposed on the anatomical MRI, and optional interactive masking is made available. After *OspreyLoad*, *OspreyProc*, and *OspreyFit*, the MRSI grid is superimposed on the anatomical MRI and accompanied by the MR spectrum at the respective analysis stage, allowing the user to visualize different voxels via click interactions. After *OspreyOverview*, an interactive atlas visualization is created, allowing the user to apply different thresholds for fractional volume, FWHM, SNR, CRLB, and SD filters and to visualize quantifications for all metabolites and anatomical regions.

#### HTML reports

Interactive HTML reports are generated in the final stage of the Osprey MRSI workflow using the Plotly graphing libraries (version 3.1.0 from https://github.com/plotly/plotly_matlab). Semi-interactive visualizations for each module are summarized in a single HTML report, enabling simple interactions, such as zooming into different regions of a spectrum or metabolic image. The HTML report can be easily shared and viewed by collaborators, as it can be opened in any web browser or integrated into larger visualization or database systems, e.g., as has been done for the Osprey outputs in the HBCD study^23^ using the LORIS database system^50^.

#### Interface with FSLeyes NIfTI Viewer

The NIfTI and NIfTI-MRS files of Osprey-MRSI can be interactively visualized in FSLeyes (version 0.1.17.0) using the mrs-plugin (version 0.1.7), which supports defining arbitrary derivative file structures via a file tree. This file tree includes the naming logic for Osprey-MRSI outputs and a default color scheme, both of which Osprey-MRSI generates automatically. This enables straightforward visualization with other MR images or derivatives from other pipelines.

### Workflow demonstration

Various 3T MRSI datasets were processed to demonstrate the versatility of Osprey-MRSI, including *in vitro* and *in vivo* MRSI data from different vendors, sites, and MRSI sequences. The analysis was performed using MATLAB R2023a.

#### In vitro MRSI experiments

*In vitro* MRSI experiments were performed at 3T for three major MR vendors (GE, Philips, Siemens) to validate the correct coregistration of the MRSI datasets and the MR images. Experimental details can be found in **Supplementary Material 2**. The coregistration was confirmed by visualizing the MRSI water signals on the MR images and comparing the MRSI spectra from different locations in Osprey and FSLeyes.

#### In vivo MRSI experiments

*In vivo* MRSI experiments were performed for three major MR vendors. Two Philips *in vivo* 2D multi-slice spin-echo MRSI datasets^51^ from healthy volunteers were analyzed, including a medium-TE (70 ms) and a short-TE (15 ms) MRSI. For GE, single-slice sLASER^52^ MRSI datasets from a healthy volunteer at 30 and 70 ms TE were analyzed. For Siemens, single-slice PRESS MRSI datasets at 35 and 280 ms TE from a clinical consult were analyzed. The study was conducted in accordance with local regulatory requirements. Experimental details and subject information are provided in **Supplementary Materials 3** and **4**, respectively. Basis sets matching the vendor-specific sequence timings and RF pulse shapes were simulated using *MRSCloud*^53^. The underlying preprocessing steps and model procedures were adapted to reflect the experimental conditions and demonstrate different aspects of the workflow (for details, see the minimum reporting standards table in **Supplementary Material 5**). To assess the benefits of parallelization, modeling of the Philips TE 70 ms was performed on two different machines with the following specifications: 2.4 GHz processor with 4 logical cores and 8 GB memory (Machine A) and 5.5 GHz processor with 24 logical cores and 192 GB memory (Machine B). Modeling on machine A was also performed without parallelization.

## Results

All MRSI datasets were successfully analyzed in Osprey-MRSI.

### In vitro MRSI experiments

The correct coregistration of MRSI data and MR images was confirmed for the *in vitro* datasets from Philips, GE, and Siemens (**Supplementary Material 6**). Both Osprey-MRSI and FSLeyes show the same localization and spectra for all example spectra.

### In vivo MRSI experiments

**Figure 2** summarizes representative *in vivo* results from the Philips medium-TE MRSI dataset. Automated brain and lipid masks and tissue fraction maps at MRSI resolution are overlaid on the T_1_-weighted image (**Figure 2A**), confirming the coregistration between MRSI data and MR images. The processed spectra and corresponding LCM results from two voxels are visualized alongside the voxel localization and the quality filter map (**Figure 2B**). Fully tissue- and relaxation-corrected metabolic images and relative CRLB images for tNAA, tCr, and tCho indicate successful LCM across the whole slice (**Figure 2C**) with concentration and CRLB estimates are consistent with literature. Similar observations are shown for the atlas-based analysis in the bilateral thalamus for the same metabolites (**Figure 2D**). The global concentration estimates for tCho indicate higher choline concentrations in white matter than in gray matter, as previously reported (**Figure 2E**)^8,54,55^. The time to model all voxels from the metabolite dataset was 19 minutes 29 seconds on Machine A without parallelization, 8 minutes 24 seconds with parallelization, and 2 minutes on Machine B, clearly show the benefits of fully implemented parallelization and modern hardware.

**Figure 2.**
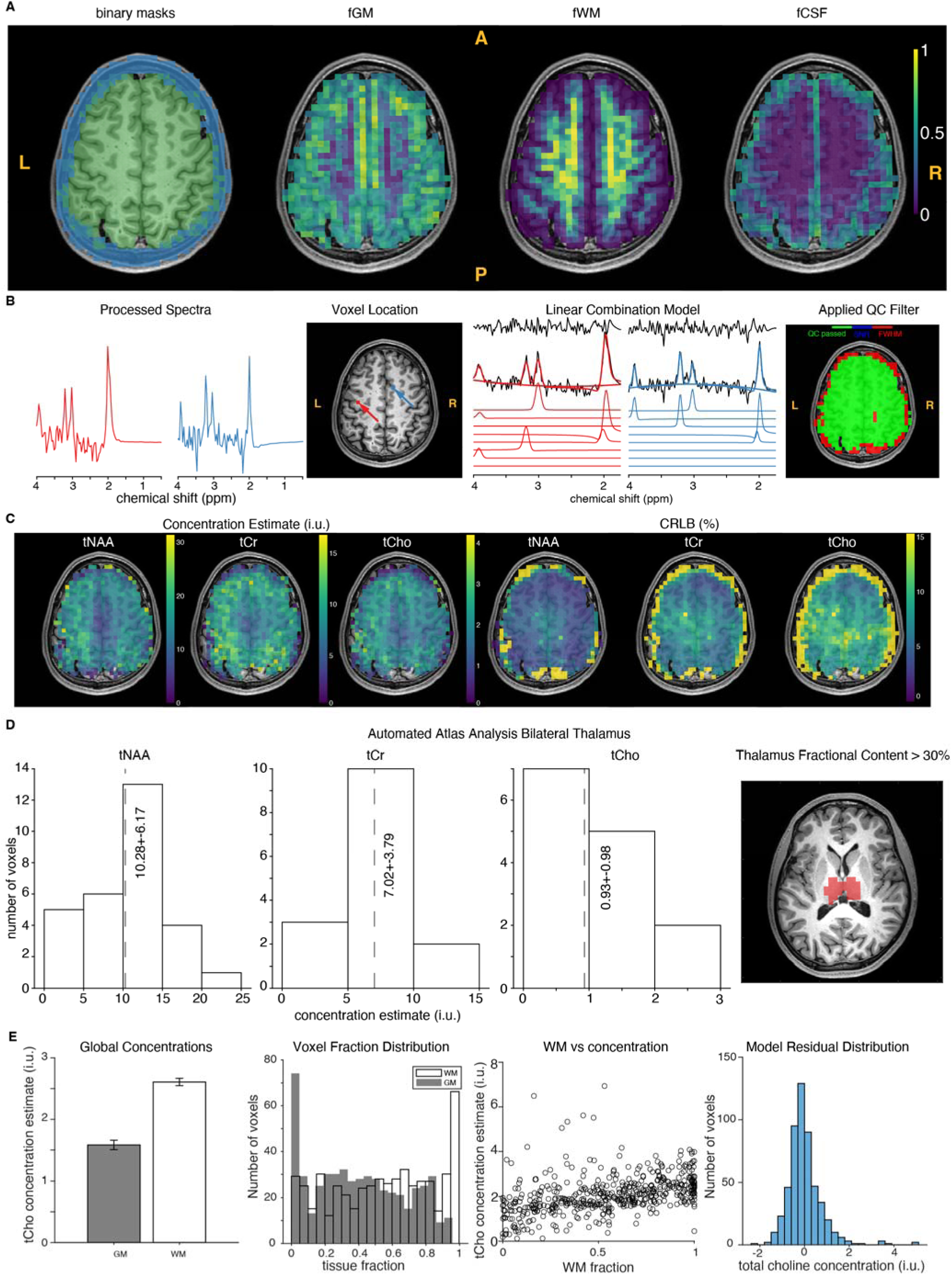
Example Osprey workflow results from the medium-TE MRSI dataset. (A) Visualization of binary lipid and brain masks as well as tissue fraction maps (only one MRSI slice shown for clarity). (B) Example of processed spectra, linear-combination model results, and quality control. Voxel locations from the example spectra are shown on the T1-weighted image. (C) Fully tissue- and relaxation-corrected metabolic maps and relative CRLB maps for tNAA, tCr, and tCho from a representative MRSI slice. Note that the presented data in panels A-C are from the same slice (overlayed on the same anatomical image). (D) Example atlas analysis from the bilateral thalamus region with distributions for fully tissue- and relaxation-corrected estimates of tNAA, tCr, and tCho. (E) Global concentration analysis for tCho showing global concentration estimates after linear regression, voxel fraction distribution, linear regression analysis, and model residuals.

**Figure 3** summarizes *in vivo* results of the Philips short-TE MRSI dataset. The coregistration of the MRSI data and T_1_-weighted image is visualized across all 5 MRSI slices (**Figure 3A**). LCM results from four example voxels are visualized alongside the voxel localization (**Figure 3B**). Fully tissue-and-relaxation corrected metabolic images and relative CRLB images from two example slices are shown alongside an atlas-based analysis of the bilateral thalamus for tNAA, tCr, tCho, mI, and Glx (**Figure 3C**). Again, the results agree well with previous literature reports^8,54,55^.

**Figure 3.**
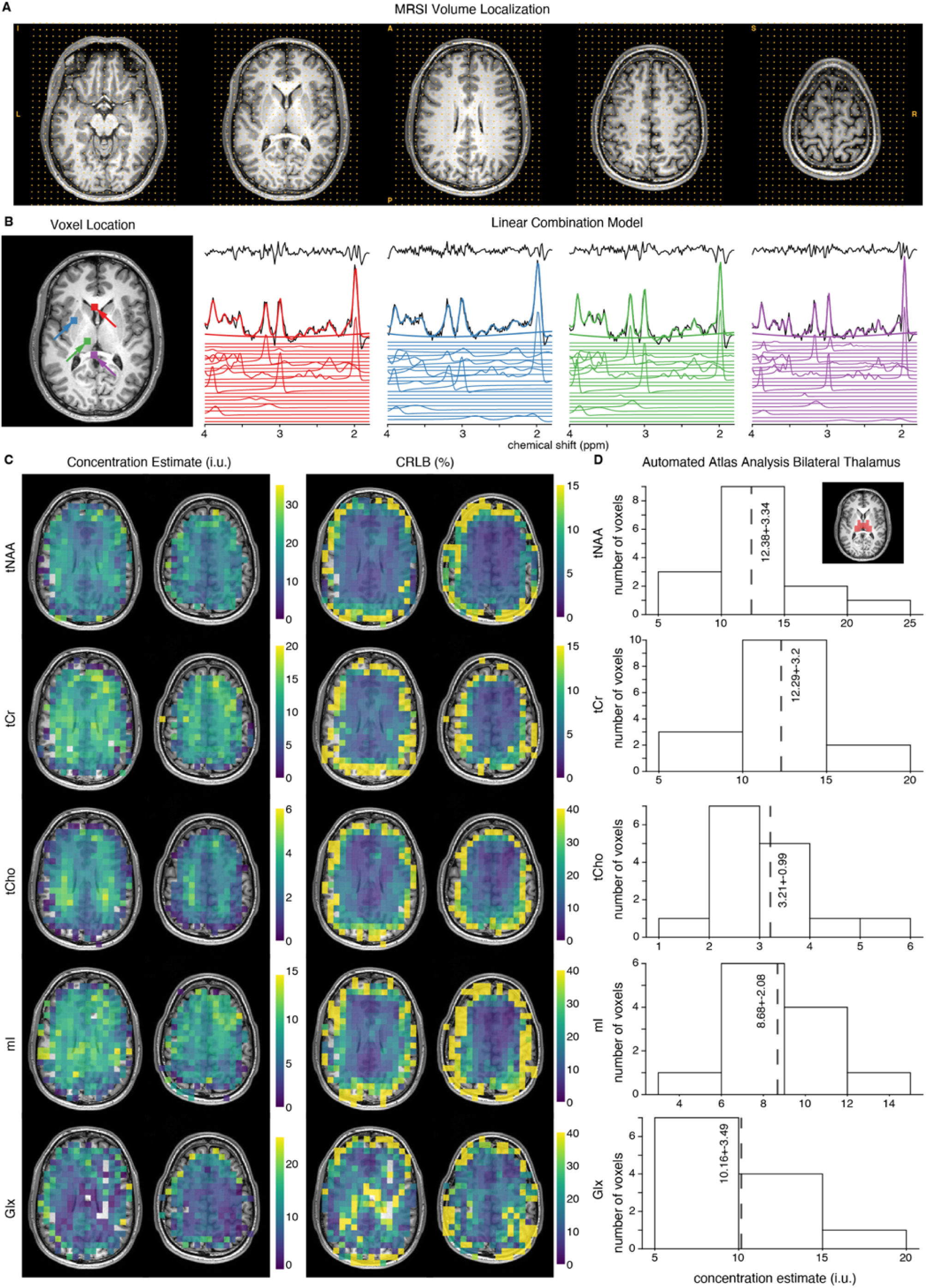
Example Osprey workflow results from the short-TE MRSI dataset. (A) Visualization of the MRSI voxels across all 5 slices. (B) Example of linear-combination model results. Voxel locations from the example spectra are shown on the T**1**-weighted image. (C) Two representative slices with fully tissue- and relaxation-corrected metabolic maps and relative CRLB maps (D) tNAA, tCr, tCho, mI, and Glx distributions from the bilateral thalamus as example location.

Successful analysis results for the GE and Siemens datasets are not shown in the manuscript for clarity, but are available in **Supplementary Materials 7** and **8**.

### Interactive Visualization

***Figure 4*** shows the three interactive visualizations in Osprey-MRSI, including an interactive data viewer for inspecting MRSI spectra at each step of the analysis (***Figure 4A***). Interactive masking (***Figure 4B***) allows the user to manually define an analysis. The interactive atlas analysis interface (***Figure 4C***) allows the user to inspect different regions of interest and metabolite quantifications, and to apply various quality-control filters. Static examples of the HTML report are shown in Supplementary Material 9; interactive examples are available on the Open Science Framework^26^ and the Osprey-MRSI GitHub repository.

**Figure 4.**
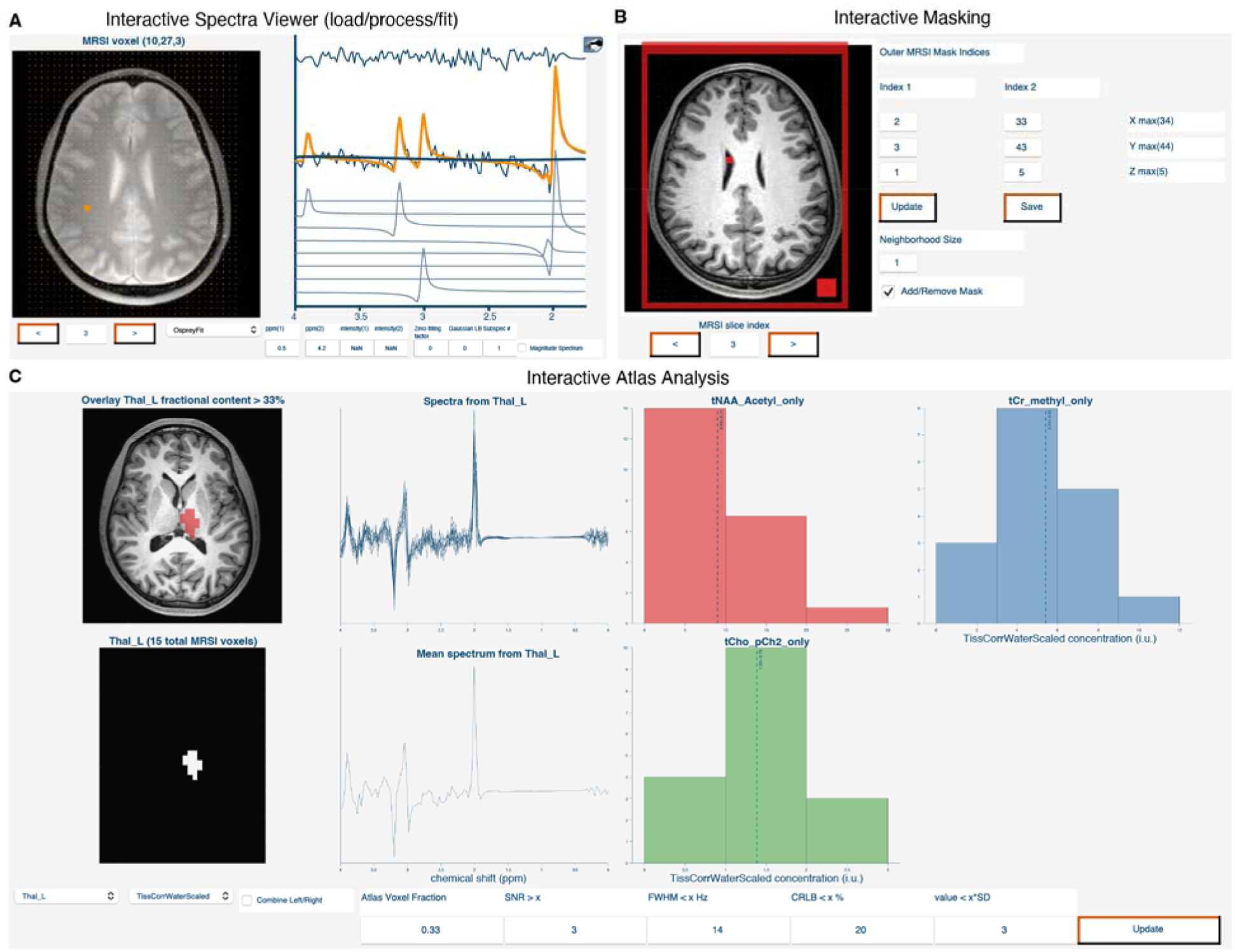
Interactive visualization in Osprey. (A) MRSI data visualization to inspect data after loading, processing or modeling. The left side of the window shows the user-supplied image (T1-weighted anatomical or MRSI localizer) and the current MRSI voxel. The right side shows the MR spectrum or LCM results. The MRSI slice, axis limits, zero-filling or linebroadening, real/magnitude or subspectra can be changed on the fly. (B) Interactive masking can be used to manually define voxels of interest. This can be used to overwrite the automated brain mask or in vitro data analysis. (C) Interactive atlas analysis. Visualizes MRSI spectra and metabolite distributions from the atlas region. The user can change the atlas region, combine left/right, and change the quality control thresholds on the fly.

**Figure 5** visualizes full compatibility between Osprey derivatives and FSLeyes via the file tree.

**Figure 5.**
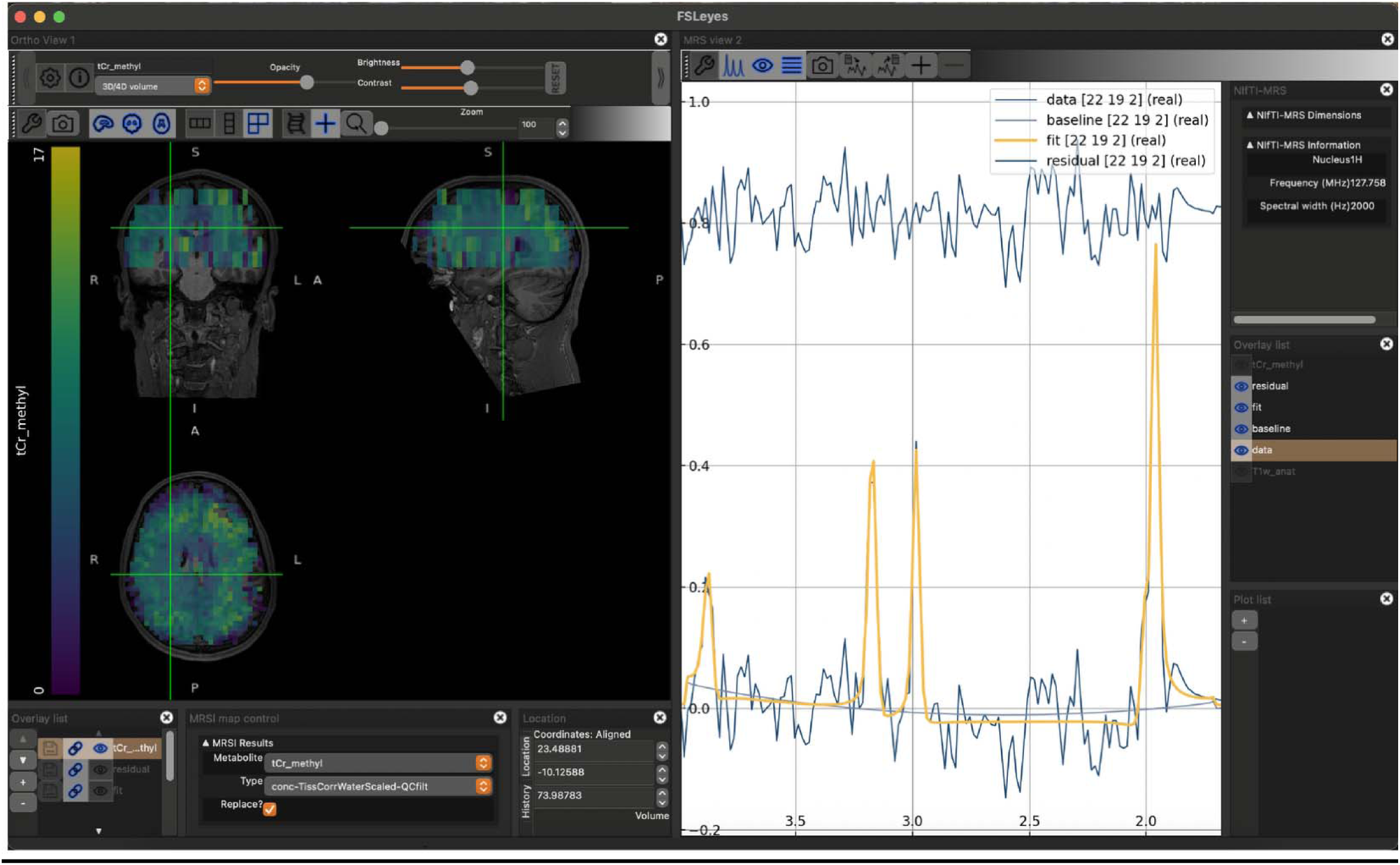
Example of interactive visualization of Osprey workflow results in FSLeyes. The MRSI map naming conventions were automatically parsed from the file tree and are represented in the MRSI map control panel.

## Discussion

Even though ^1^H-MRSI is a versatile tool for spatially resolving *in vivo* metabolism, end-to-end analysis of ^1^H- MRSI is challenging due to high dimensionality, spectral overlap, and heterogeneous spectral quality across the MRSI volume. Fully automated analysis pipelines are best suited to overcome this challenge; however, only a few software solutions exist for end-to-end ^1^H-MRSI data analysis. In this study, we presented a newly implemented ^1^H-MRSI workflow Osprey-MRSI, a modular MATLAB-based open-source ^1^H-MRS analysis package.

Osprey-MRSI features include spatial transformation and filtering, automated brain and lipid masking, a modular linear-combination algorithm with explicit B_0_ frequency-shift correction, and fully tissue- and relaxation-corrected quantification. To make the high-dimensional MRSI results more digestible, atlas-based statistics and global concentrations based on linear regression are also calculated. Finally, Osprey-MRSI provides interactive visualization of the MRSI data and analysis results and interfaces with FSLeyes via NIfTI-MRS.

Osprey-MRSI is the first MATLAB-based self-contained end-to-end ^1^H-MRSI analysis pipeline for MRSI data from clinical MR scanners. Previous MATLAB-based workflows relied on LCModel as an external model algorithm^3,4,12^. The novel generalized LCM algorithm enables flexible implementation of model procedures, which is a major advantage of Osprey-MRSI, as demonstrated in this study by using explicit B_0_ frequency-shift correction and two different baseline approaches for short- and medium-TE MRSI data modeling, respectively. Thus, the algorithm enables researchers to investigate the impact of the modeling meta-parameter on metabolite estimation in MRSI data much more easily than previously possible by modifying the *model procedure* file. However, because each *model procedure* file is self-contained, reproducibility and comparability are ensured, as it can be distributed with the results for replication. Similarly, each module in the workflow can be modified to investigate the impact of each processing stage on metabolite estimation. Osprey-MRSI offers features similar to other self-contained open-source ^1^H-MRSI workflows, such as MIDAS^2^, FSL-MRS^6^, and ABFit^10,11^, and provides a MATLAB-based alternative, which is a valuable addition, based on our experience with Osprey for single-voxel MRS. The integration of standardized brain atlases and automated quality-control thresholding is of particular interest for application-oriented MRSI research, fostering group-based statistics and multimodal analysis of the same brain regions. It also allows generating population-based metabolic maps in standard space, like MIDAS, for direct comparison of single-subject results with the population. Further data inspection is enabled by interfacing with FSLeyes through the NIfTI-MRS community standard and file tree descriptions, allowing for coregistered visualization of the MRSI analysis results and other imaging modalities.

For this study, we focused on the workflow demonstration and therefore only included a limited number of short- and medium-TE MRSI datasets from all three major MRI vendors. Earlier versions of this workflow have been used in larger study populations to assess the reproducibility of short-TE and GABA-edited multi-slice MRSI^56^ and to detect metabolic alterations in individuals with tuberous sclerosis complex^57^. Future studies are needed to benchmark the workflow against existing MRSI analysis workflows, for example, by comparing it to the recent ISMRM MRS Study Group Fitting Challenge results on MRSI data^58^. However, the analysis results from the workflow demonstration presented in this study agree well with previous literature, including the range of concentration estimates and CRLBs. Similarly, spatial distributions reflect common literature findings, such as increased total choline in white matter compared to gray matter and in frontal brain regions. Compared to other MRSI analysis workflows^15^, Osprey-MRSI does not directly support DICOM export or PACS integration. However, Osprey has previously been compiled^24^ and used in a large multi-center study^23^, with a LORIS database to integrate all derivatives^50^. Thus, offering a scalable solution for centralized processing on high-performance systems to fully leverage Osprey-MRSI’s computational parallelization.

## Conclusion

Osprey-MRSI offers state-of-the-art methods with minimal user interaction, making it accessible to non-expert users. This workflow will increase access to standardized MRSI analysis for application-oriented MRSI research. Osprey-MRSI’s modularity, open-source environment, and the generalized modeling algorithm will foster innovation and development of novel MRSI-specific analysis methods.

## Supporting information

Data Availability Statement

## Acknowledgement

The authors thank Dr. Ipek Ozdemir (UC Davis) for ongoing discussions during our weekly group meeting.

This work has been supported in part by NIH grants R01EB028259, R01NS134694, R00AG062230, R21EB033516, K99/R00 AG080084, and P41EB031771, as well as by DOD grant W81XWH2010819. WTC is supported by Wellcome [225924/Z/22/Z]. This research was funded in whole, or in part, by the Wellcome Trust. For the purpose of open access, the author has applied a CC BY public copyright license to any Author Accepted Manuscript version arising from this submission.

