## Supplementary Material for "Fully automated open-source analysis and interactive visualization of magnetic resonance spectroscopic imaging (MRSI) data in Osprey-MRSI"

***Supplementary Material 1****. Overview of end-to-end MRSI analysis pipelines*

| **Name** | **Preprocessing Features** | **Modeling  Approach** | **Other  Features** | **Code** | **Programing**  **Language** | **Published** |
| --- | --- | --- | --- | --- | --- | --- |
| MIDAS | coil-combination, spatial fourier transformation, brain masking, frequency correction, ECC, water & lipid removal | Linear-combination modeling | Automated quality control, fractional volume correction, spatial normalization, atlas analysis, coregistered GUI visualization, reports | open | IDL | 2006 |
| FSL-MRS | coil-combination, brain masking, frequency correction, ECC, water & lipid removal | Linear-combination modeling | Fractional volume correction, coregistered GUI visualization, reports, NIfTI-MRS outputs | open | python | 2018 |
| Tarquin | manual masking, frequency correction, ECC, water & lipid removal | Linear-combination modeling | coregistered GUI visualization, tabulated output | open | C++ | 2011 |
| Spant/ABFit | manual masking, coil-combination, hanning filtering, k-space zero-filling, ECC, water removal, automated phase correction | Linear-combination modeling | PG k-space extrapolation, fractional volume correction, spectral decomposition | open | R | 2021 |
| FID-A on FIRE | coil-combination, spatial fourier transformation, frequency correction, ECC, water & lipid removal | LCModel wrapper | non-cartesian reconstruction, works on Siemens scanner | open | Matlab | 2024 |
| VDI | parse external masks, automated phasing, ECC, frequency correction | LCModel wrapper | Automated quality control, fractional volume correction, spatial normalization, atlas analysis, global concentration, reports | open | Matlab | 2023 |
| MRSSpecLAB | Coil-combination, hanning filtering, ECC, water removal, automated phase correction, line broadening | LCModel wrapper | Static GUI visualization | open | python | 2025 |
| Oryx-MRSI | Frequency correction, ECC (both performed in LCModel) | LCModel wrapper | Automated quality control, chemical shift correction, fractional volume correction, spatial normalization, atlas analysis, coregistered GUI visualization, reports | open | Matlab | 2022 |
| MRS4Brain  (pre-clincal) | brain masking, ECC, water & lipid removal | LCModel wrapper | Semi-automated quality control, GUI visualization | open | Matlab | 2025 |
| jMRUI | Coil-combination, hanning filtering, ECC, water removal, automated phase correction, line broadening | Various approaches | Coregistered GUI visualization | closed | JAVA | 2009 |
| SIVIC | coil-combination, spatial fourier transformation, baseline removal, apodization, zero-filling | Peak height and peak integration | Coregistered GUI visualization, PACS integration | open | C++ | 2013 |
| LCModel | Coil-combination, frequency correction, ECC | Linear-combination modeling | Static grid output, tabulated output | open | Fortran | 1992 |
| CSX | Bandpass filter | Peak integration | Coregistered GUI visualization | closed | C++ | 2005 |

***Supplementary Material 2****. Experimental details of in vitro MRSI example datasets.*

| **Parameter** | **Philips  braino** | **GE  structural** | **Siemens  structural** |
| --- | --- | --- | --- |
| Matrix size | 22 x 28 x 5 | 8 x 8 x 1 | 16 x 16 x 1 |
| Nominal voxel size (mm) | 8 x 8 x 14 = 0.9 ml  (4 mm gap) | 30 x 30 x 10 = 9 ml | 15 x 15 x 10 = 2.25 ml |
| TR/TE | 3.5 s / 15 ms | 2 s / 30 ms | 2 s / 40 ms |

***Supplementary Material 3****. Experimental details of in vivo MRSI example datasets*

| **Vendor** | **Matrix size** | **Nominal  voxel size** | **TR / TE / averages** | **Sequence** |
| --- | --- | --- | --- | --- |
| Philips | 34 x 44 x 5 | 5 x 5 x 12 mm = 0.3 ml  (4 mm gap) | 3.5 s / 70 ms / 1  (water MRSI 2 s / 15 ms) | 2D multi slice spin-echo |
| Philips | 22 x 28 x 5 | 8 x 8 x 14 mm = 0.9 ml  (4 mm gap) | 3.5 s / 15 ms / 1 (water MRSI 2 s / 15 ms) | 2D multi slice spin-echo |
| GE | 16 x 16 x 1 | 15 x 15 x 20 mm = 4.5 ml | 2 s / 30 ms / 1 | 2D sLASER |
| GE | 16 x 16 x 1 | 15 x 15 x 20 mm = 4.5 ml | 2 s / 70 ms / 1 | 2D sLASER |
| Siemens | 16 x 16 x 1 | 10 x 10 x 15 mm = 1.5 ml | 1.7 s / 35 ms / 1 | 2D PRESS |
| Siemens | 16 x 16 x 1 | 10 x 10 x 15 mm = 1.5 ml | 1.7 s / 280 ms / 1 | 2D PRESS |

***Supplementary Material 4****. Subject information of in vivo MRSI example datasets. *Subject age of the GE example dataset from a healthy adult cannot be shared following local regulatory requirements.*

| **Subject** | **Age**  **(years)** | **gender** | **other comments** |
| --- | --- | --- | --- |
| Philips (medium TE) | 23 | male | healthy subject |
| Philips (short TE) | 35 | male | healthy subject |
| GE (short / medium TE | X* | female | healthy subject |
| Siemens (short / long TE) | 10 | female | clinical consult |

***Supplementary Material 5****. Summary following minimum reporting standards in MRS for in vivo MRSI example datasets. See Lin et al. ‘Minimum Reporting Standards for in vivo Magnetic Resonance Spectroscopy (MRSinMRS): Expert consensus recommendations. NMR in Biomedicine. 2012:e4484.* [doi.org/10.1002/nbm.4484](https://doi.org/10.1002/nbm.4484)

| **1. Hardware** | |
| --- | --- |
| **Medium TE Philips** |  |
| a. Field strength [T] | 3 |
| b. Manufacturer | Philips |
| c. Model (software version if available) | dStream Achieva (R5.7.1) |
| d. RF coils: nuclei (transmit/receive), number of channels, type, body part | 1H, 32, matrix coil, head |
| e. Additional hardware | - |
| **Short TE Philips** |  |
| a. Field strength [T] | 3 |
| b. Manufacturer | Philips |
| c. Model (software version if available) | dStream Ingenia Elition (R5.7.1 R2D2) |
| d. RF coils: nuclei (transmit/receive), number of channels, type, body part | 1H, 32, matrix coil, head |
| e. Additional hardware | - |
| **short TE GE** |  |
| a. Field strength [T] | 3 |
| b. Manufacturer | GE |
| c. Model (software version if available) | MR 750 (MR30.1 R01) |
| d. RF coils: nuclei (transmit/receive), number of channels, type, body part | 1H, 32, transmit array coil, head |
| e. Additional hardware | - |
| **Medium TE GE** |  |
| a. Field strength [T] | 3 |
| b. Manufacturer | GE |
| c. Model (software version if available) | MR 750 (MR30.1 R01) |
| d. RF coils: nuclei (transmit/receive), number of channels, type, body part | 1H, 32, matrix coil, head |
| e. Additional hardware | - |
| **short TE Siemens** |  |
| a. Field strength [T] | 3 |
| b. Manufacturer | Siemens |
| c. Model (software version if available) | Magentom Skyra (VE 11) |
| d. RF coils: nuclei (transmit/receive), number of channels, type, body part | 1H, 20, matrix coil, head |
| e. Additional hardware | - |
| **long TE Siemens** |  |
| a. Field strength [T] | 3 |
| b. Manufacturer | Siemens |
| c. Model (software version if available) | Magentom Skyra (VE 11) |
| d. RF coils: nuclei (transmit/receive), number of channels, type, body part | 1H, 20, matrix coil, head |
| e. Additional hardware | - |

| **2. Acquisition** | |
| --- | --- |
| **Medium TE Philips** | |
| a. Pulse sequence | 2D spin-echo multi-slice (Duyn et al. 1993) |
| b. Volume of interest (VOI) locations | Example automated atlas analysis of the Pallidum |
| c. Nominal VOI size LR x AP x HF [mm3] | 5 x 5 x 12 |
| d. Repetition time (TR), echo time (TE) [ms] | TR 3500 ms, TE 70 ms |
| e. Total number of averages per spectrum | 1 total averages |
| f. Additional sequence parameters | F1: 2000 Hz, 512 points 2D: 170 x 220 x 76 mm3 FOV, 4 mm gap, matrix size: 34 x 44 x 5, elliptical k-space sampling |
| g. Water suppression method | Hypergeometric dual-band (water and lipid) + 8 OVS (Zhu et al. 2010) |
| h. Shimming method, reference peak, and threshold of acceptance of shim chosen | second order pencil-beam volume, water, - |
| i. Trigger or motion correction | - |
| **Short TE Philips** | |
| a. Pulse sequence | 2D spin-echo multi-slice (Duyn et al. 1993) |
| b. Volume of interest (VOI) locations | Example automated atlas analysis of the Pallidum |
| c. Nominal VOI size LR x AP x HF [mm3] | 8 x 8 x 14 |
| d. Repetition time (TR), echo time (TE) [ms] | TR 3500 ms, TE 15 ms |
| e. Total number of averages per spectrum | 1 total averages |
| f. Additional sequence parameters | F1: 2000 Hz, 512 points 2D: 176 x 224 x 86 mm3 FOV, 4 mm gap, matrix size: 22 x 28 x 5, elliptical k-space sampling |
| g. Water suppression method | Hypergeometric dual-band (water and lipid) + 8 OVS (Zhu et al. 2010) |
| h. Shimming method, reference peak, and threshold of acceptance of shim chosen | second order pencil-beam volume, water, - |
| i. Trigger or motion correction | - |
| **Short TE GE** | |
| a. Pulse sequence | 2D sLASER MRSI (adapted Deelchand et al. 2021) |
| b. Volume of interest (VOI) locations | Example automated atlas analysis of the Pallidum |
| c. Nominal VOI size LR x AP x HF [mm3] | 15 x 15 x 20 |
| d. Repetition time (TR), echo time (TE) [ms] | TR 2000 ms, TE 30 ms |
| e. Total number of averages per spectrum | 1 total averages |
| f. Additional sequence parameters | F1: 5000 Hz, 4096 points 2D: 240 x 240 x 20 mm3 FOV, matrix size: 16 x 16 x 1 |
| g. Water suppression method | VAPOR with OVS |
| h. Shimming method, reference peak, and threshold of acceptance of shim chosen | Dual-echo GRE (higher order for in-vivo) |
| i. Trigger or motion correction | - |
| **Medium TE GE** | |
| a. Pulse sequence | 2D sLASER MRSI (adapted Deelchand et al. 2021) |
| b. Volume of interest (VOI) locations | Example automated atlas analysis of the Pallidum |
| c. Nominal VOI size LR x AP x HF [mm3] | 15 x 15 x 20 |
| d. Repetition time (TR), echo time (TE) [ms] | TR 2000 ms, TE 70 ms |
| e. Total number of averages per spectrum | 1 total averages |
| f. Additional sequence parameters | F1: 5000 Hz, 4096 points 2D: 240 x 240 x 20 mm3 FOV, matrix size: 16 x 16 x 1 |
| g. Water suppression method | VAPOR with OVS |
| h. Shimming method, reference peak, and threshold of acceptance of shim chosen | Dual-echo GRE, higher order shim |
| i. Trigger or motion correction | - |
| **short TE Siemens** | |
| a. Pulse sequence | 2D PRESS MRSI (Bottomley 1985) |
| b. Volume of interest (VOI) locations | Brain lesion + normal appearing brain |
| c. Nominal VOI size LR x AP x HF [mm3] | 10 x 10 x 15 |
| d. Repetition time (TR), echo time (TE) [ms] | TR 1700 ms, TE 35 ms |
| e. Total number of averages per spectrum | 1 total averages |
| f. Additional sequence parameters | F1: 2000 Hz, 512 points 2D: 160 x 160 x 15 mm3 FOV, 2.5 mm gap, matrix size: 16 x 16 x 1 |
| g. Water suppression method | CHESS |
| h. Shimming method, reference peak, and threshold of acceptance of shim chosen | Automated brain shim |
| i. Trigger or motion correction | - |
| **long TE Siemens** | |
| a. Pulse sequence | 2D PRESS MRSI (Bottomley 1985) |
| b. Volume of interest (VOI) locations | Brain lesion + normal appearing brain |
| c. Nominal VOI size LR x AP x HF [mm3] | 10 x 10 x 15 |
| d. Repetition time (TR), echo time (TE) [ms] | TR 1700 ms, TE 280 ms |
| e. Total number of averages per spectrum | 1 total averages |
| f. Additional sequence parameters | F1: 2000 Hz, 512 points 2D: 160 x 160 x 15 mm3 FOV, matrix size: 16 x 16 x 1 |
| g. Water suppression method | CHESS |
| h. Shimming method, reference peak, and threshold of acceptance of shim chosen | Automated brain shim |
| i. Trigger or motion correction | - |

| **3. Data analysis methods and outputs** | |
| --- | --- |
| **Medium TE Philips** | |
| a. Analysis software | Osprey-MRSI 1.0.0 (MRSI-Release) code available on Osprey-MRSI release page; Modeling with gLCM algorithm (Zöllner et al. 2024) |
| b. Processing steps settings | First-point FID phase correction; L2-basis water and lipid removal; cross-correlation frequency correction with baseline removal; |
| c. Output measure | Fully tissue- and relaxation-corrected metabolite estimate maps (Gasparovic et al. 2006) |
| d. Quantification references and assumptions, fitting model assumptions | Basis set list: Cr_methyl, Cr_metylene, GPC_pCh2, NAA_Acetyl, NAAG_Acetyl, PCh_trimethyl, PCr_ch2nhnh, PCr_ch3 Fitting method: GeneralizedBasicPhysicsModelWithGlobalFreqShift.m (Simicic et al. 2025) frequency domain model 1.75 to 4 ppm with second order polynomial baseline. NAA_Acetyl, NAAG_Acetyl soft constraint. See procedures/1StepFinal_in_vivo_longTE.json for details. |
| **Short TE Philips** | |
| a. Analysis software | Osprey-MRSI 1.0.0 (MRSI-Release) code available on Osprey-MRSI release page; Modeling with gLCM algorithm (Zöllner et al. 2024) |
| b. Processing steps settings | First-point FID phase correction; HSVD water removal; cross-correlation frequency correction with baseline removal; |
| c. Output measure | Fully tissue- and relaxation-corrected concentration estimates (Gasparovic et al. 2006) |
| d. Quantification references and assumptions, fitting model assumptions | Basis set list: Asc, Asp, Cr, CrCH2, GABA, GPC, GSH, Gln, Glu, mI, Lac, NAA, NAAG, PCh, PCr, PE, sI, Tau MM basis functions: MM2.03, MM3.1, MM3.7, MM3.8, MM4.0, Lip2.0 soft constraints relative to tNAA Fitting method: GeneralizedBasicPhysicsModelWithGlobalFreqShift.m (Simicic et al. 2025) frequency domain model 1.8 to 4 ppm with regularized p-spline baseline (0.0667 ppm knot spacing, 20 AIC steps with m = 14) (see Wilson et al. 2022). See procedures/ 3Step_Spline_invivo_Reg_Optim_MRSI.json for details. |
| **Short TE GE** | |
| a. Analysis software | Osprey-MRSI 1.0.0 (MRSI-Release) code available on Osprey-MRSI release page; Modeling with gLCM algorithm (Zöllner et al. 2024) |
| b. Processing steps settings | First-point FID phase correction; L2-basis water and L2-mask lipid removal; cross-correlation frequency correction with baseline removal; |
| c. Output measure | Creatine referenced metabolite maps |
| d. Quantification references and assumptions, fitting model assumptions | Basis set list: Asc, Asp, Cr, CrCH2, GABA, GPC, GSH, Gln, Glu, mI, Lac, NAA, NAAG, PCh, PCr, PE, sI, Tau MM basis functions: MM2.03, MM3.1, MM3.7, MM3.8, MM4.0, Lip2.0 soft constraints relative to tNAA Fitting method: GeneralizedBasicPhysicsModelWithGlobalFreqShift.m (Simicic et al. 2025) frequency domain model 1.8 to 4 ppm with regularized p-spline baseline (0.0667 ppm knot spacing, 20 AIC steps with m = 14) (see Wilson et al. 2022). See procedures/ 3Step_Spline_invivo_Reg_Optim_MRSI.json for details. |
| **Medium TE GE** | |
| a. Analysis software | Osprey-MRSI 1.0.0 (MRSI-Release) code available on Osprey-MRSI release page; Modeling with gLCM algorithm (Zöllner et al. 2024) |
| b. Processing steps settings | Auto phase correction; L2-basis water and lipid removal; cross-correlation frequency correction with baseline removal; |
| c. Output measure | Creatine referenced metabolite maps |
| d. Quantification references and assumptions, fitting model assumptions | Basis set list: Cr_methyl, Cr_metylene, GPC_pCh2, NAA_Acetyl, NAAG_Acetyl, PCh_trimethyl, PCr_ch2nhnh, PCr_ch3 Fitting method: GeneralizedBasicPhysicsModelWithGlobalFreqShift.m (Simicic et al. 2025) frequency domain model 1.75 to 4 ppm with second order polynomial baseline. NAA_Acetyl, NAAG_Acetyl soft constraint. See procedures/1StepFinal_in_vivo_longTE.json for details. |
| **Short TE Siemens** | |
| a. Analysis software | Osprey-MRSI 1.0.0 (MRSI-Release) code available on Osprey-MRSI release page; Modeling with gLCM algorithm (Zöllner et al. 2024) |
| b. Processing steps settings | First-point FID phase correction; L2-basis water and L2-mask lipid removal; cross-correlation frequency correction with baseline removal; |
| c. Output measure | Creatine referenced metabolite maps |
| d. Quantification references and assumptions, fitting model assumptions | Basis set list: Asc, Asp, Cr, CrCH2, GABA, GPC, GSH, Gln, Glu, mI, Lac, NAA, NAAG, PCh, PCr, PE, sI, Tau MM basis functions: MM2.03, MM3.1, MM3.7, MM3.8, MM4.0, Lip2.0 soft constraints relative to tNAA Fitting method: GeneralizedBasicPhysicsModelWithGlobalFreqShift.m (Simicic et al. 2025) frequency domain model 1.8 to 4 ppm with regularized p-spline baseline (0.0667 ppm knot spacing, 20 AIC steps with m = 14) (see Wilson et al. 2022). See procedures/ 3Step_Spline_invivo_Reg_Optim_MRSI.json for details. |
| **Long TE Siemens** | |
| a. Analysis software | Osprey-MRSI 1.0.0 (MRSI-Release) code available on Osprey-MRSI release page; Modeling with gLCM algorithm (Zöllner et al. 2024) |
| b. Processing steps settings | First-point FID phase correction; L2-basis water and lipid removal; cross-correlation frequency correction with baseline removal; |
| c. Output measure | Creatine referenced metabolite maps |
| d. Quantification references and assumptions, fitting model assumptions | Basis set list: Cr_methyl, Cr_metylene, GPC_pCh2, NAA_Acetyl, NAAG_Acetyl, PCh_trimethyl, PCr_ch2nhnh, PCr_ch3 Fitting method: GeneralizedBasicPhysicsModelWithGlobalFreqShift.m (Simicic et al. 2025) frequency domain model 1.75 to 4 ppm with second order polynomial baseline. NAA_Acetyl, NAAG_Acetyl soft constraint. See procedures/1StepFinal_in_vivo_longTE.json for details. |

| **4. Data quality** | |
| --- | --- |
| **Medium TE Philips** | |
| a. SNR (tNAA), linewidth (tNAA) [Hz] | example voxels in Figure 2: |
| b. Data exclusion criteria | visible lipid contamination or OOV echoes, automated QC thresholds: SNR > 3, Cr FWHM < 14, relative CRLB > 25%*,* Mi < 97^th^ percentile for each metabolite estimate Mi |
| c. Quality measures of postprocessing model fitting | CRLBs of target metabolites example voxels in Figure 2: |
| d. Mean and individual spectra | Example spectra in Figure 2 |
| **Short TE Philips** | |
| a. SNR (tNAA), linewidth (tNAA) [Hz] | Example voxels in Figure 3: |
| b. Data exclusion criteria | visible lipid contamination or OOV echoes, automated QC thresholds: SNR > 3, Cr FWHM < 14, relative CRLB > 25%*,* Mi < 97^th^ percentile for each metabolite estimate Mi |
| c. Quality measures of postprocessing model fitting | CRLBs of target metabolites example voxels in Figure 3: |
| d. Mean and individual spectra | Example spectra in Figure 3 |
| **Short TE GE** | |
| a. SNR (tNAA), linewidth (tNAA) [Hz] | Example voxels in Supplementary Material 7 |
| b. Data exclusion criteria | visible lipid contamination or OOV echoes, automated QC thresholds: SNR > 3, Cr FWHM < 14, relative CRLB > 25%*,* Mi < 97^th^ percentile for each metabolite estimate Mi |
| c. Quality measures of postprocessing model fitting | CRLBs of target metabolites example voxels in Supplementary Material 8: |
| d. Mean and individual spectra | Example spectra in Supplementary Material 7 |
| **Medium TE GE** | |
| a. SNR (tNAA), linewidth (tNAA) [Hz] | Example voxels in Supplementary Material 7 |
| b. Data exclusion criteria | visible lipid contamination or OOV echoes, automated QC thresholds: SNR > 3, Cr FWHM < 14, relative CRLB > 25%*,* Mi < 97^th^ percentile for each metabolite estimate Mi |
| c. Quality measures of postprocessing model fitting | CRLBs of target metabolites example voxels in Supplementary Material 8: |
| d. Mean and individual spectra | Example spectra in Supplementary Material 7 |
| **Short TE Siemens** | |
| a. SNR (tNAA), linewidth (tNAA) [Hz] | example voxels in Supplementary Material 8 |
| b. Data exclusion criteria | visible lipid contamination or OOV echoes, automated QC thresholds: SNR > 3, Cr FWHM < 14, relative CRLB > 25%*,* Mi < 97^th^ percentile for each metabolite estimate Mi |
| c. Quality measures of postprocessing model fitting | CRLBs of target metabolites example voxels in Supplementary Material 9: |
| d. Mean and individual spectra | Example spectra in Supplementary Material 8 |
| **Long TE Siemens** | |
| a. SNR (tNAA), linewidth (tNAA) [Hz] short-TE MRSI | example voxels in Supplementary Material 8 |
| b. Data exclusion criteria | visible lipid contamination or OOV echoes, automated QC thresholds: SNR > 3, Cr FWHM < 14, relative CRLB > 25%*,* Mi < 97^th^ percentile for each metabolite estimate Mi |
| c. Quality measures of postprocessing model fitting | CRLBs of target metabolites example voxels in Supplementary Material 9: |
| d. Mean and individual spectra | Example spectra in Supplementary Material 8 |

***Supplementary Material 6****.* *Visualization of in vitro MRSI examples confirming the correct coregistration in Osprey and FSLeyes for each vendor (Philips, GE, Siemens). Small differences in the images may be due to differences in image interpolation when transforming the images.*

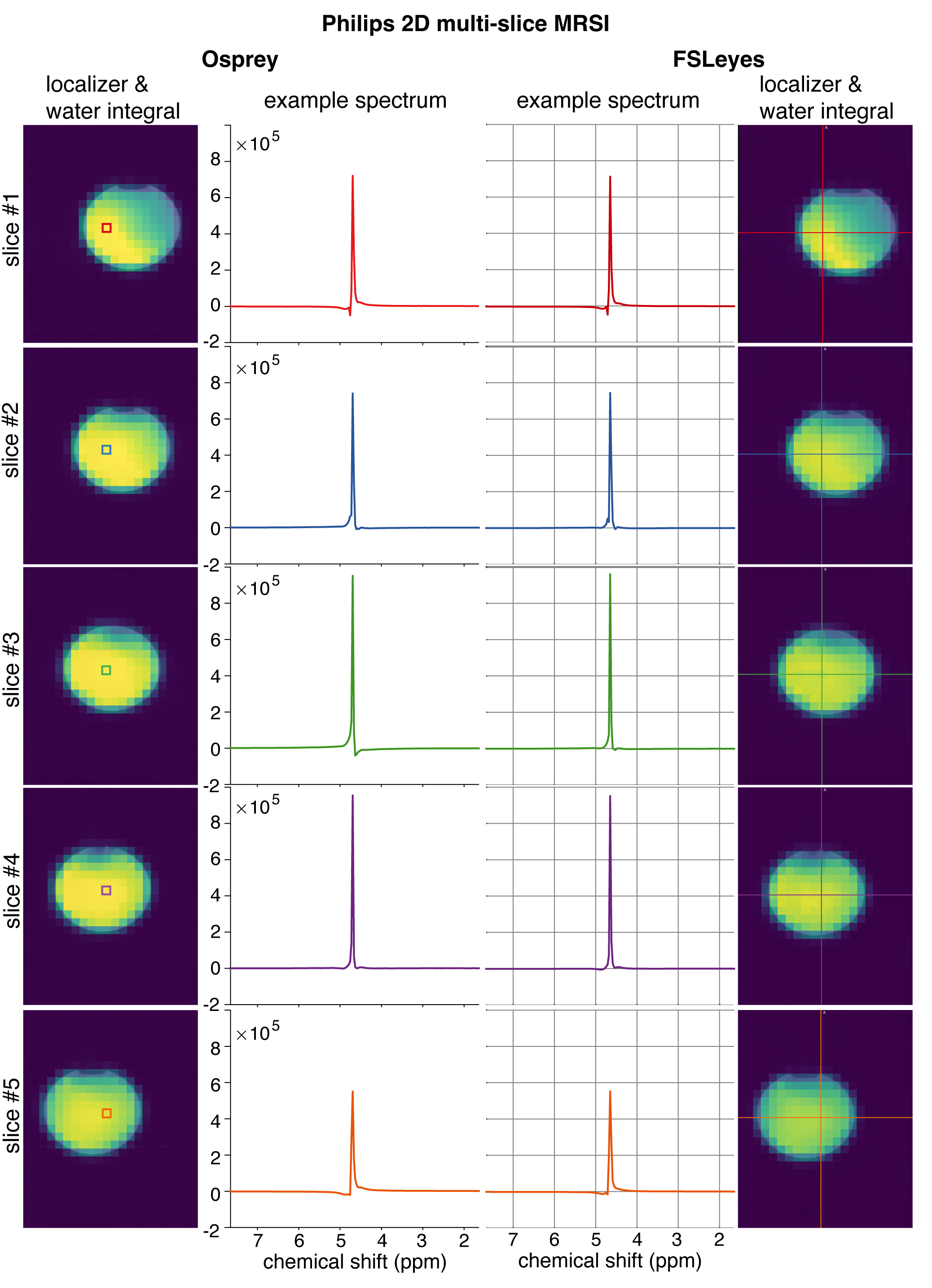

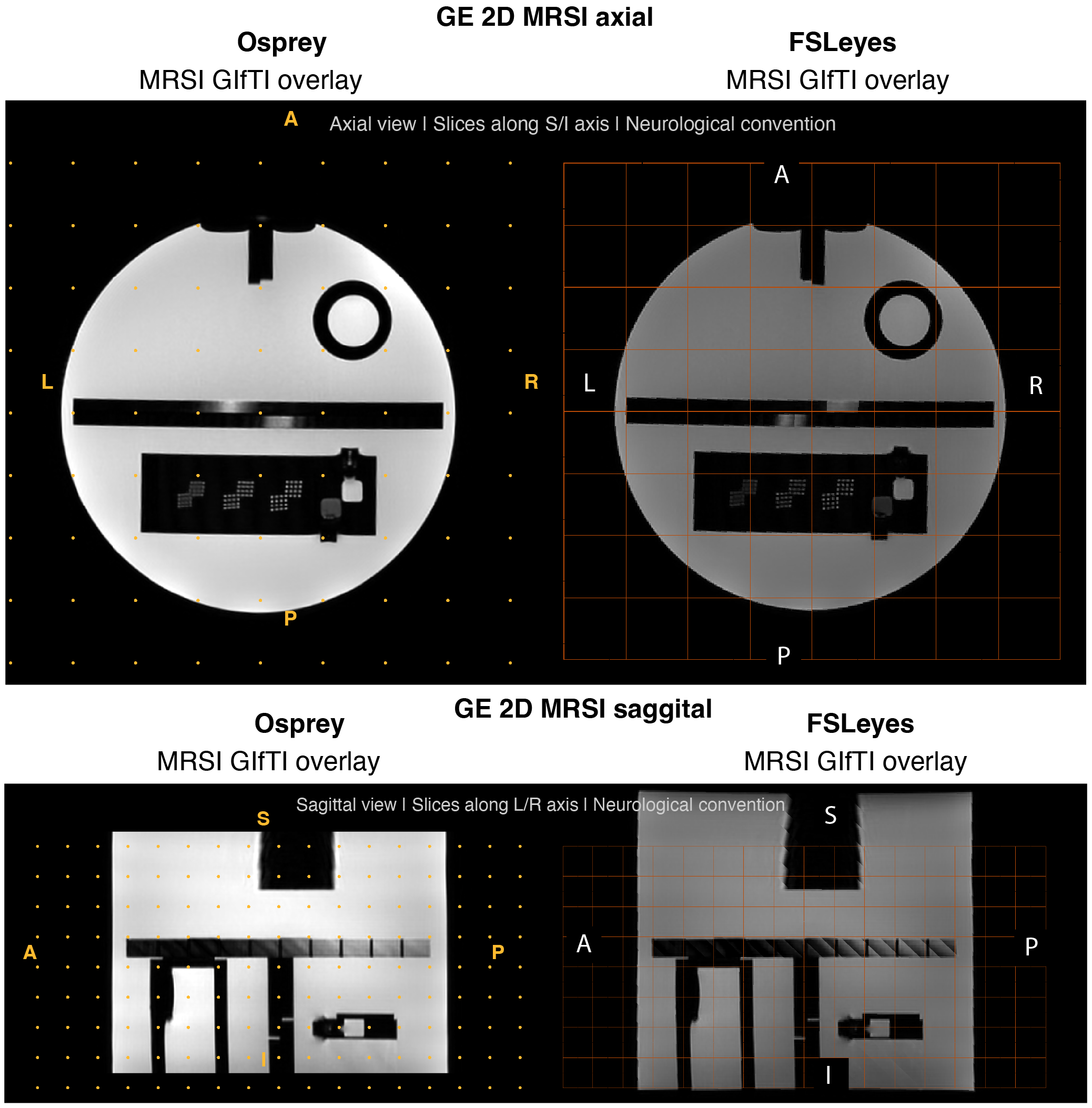

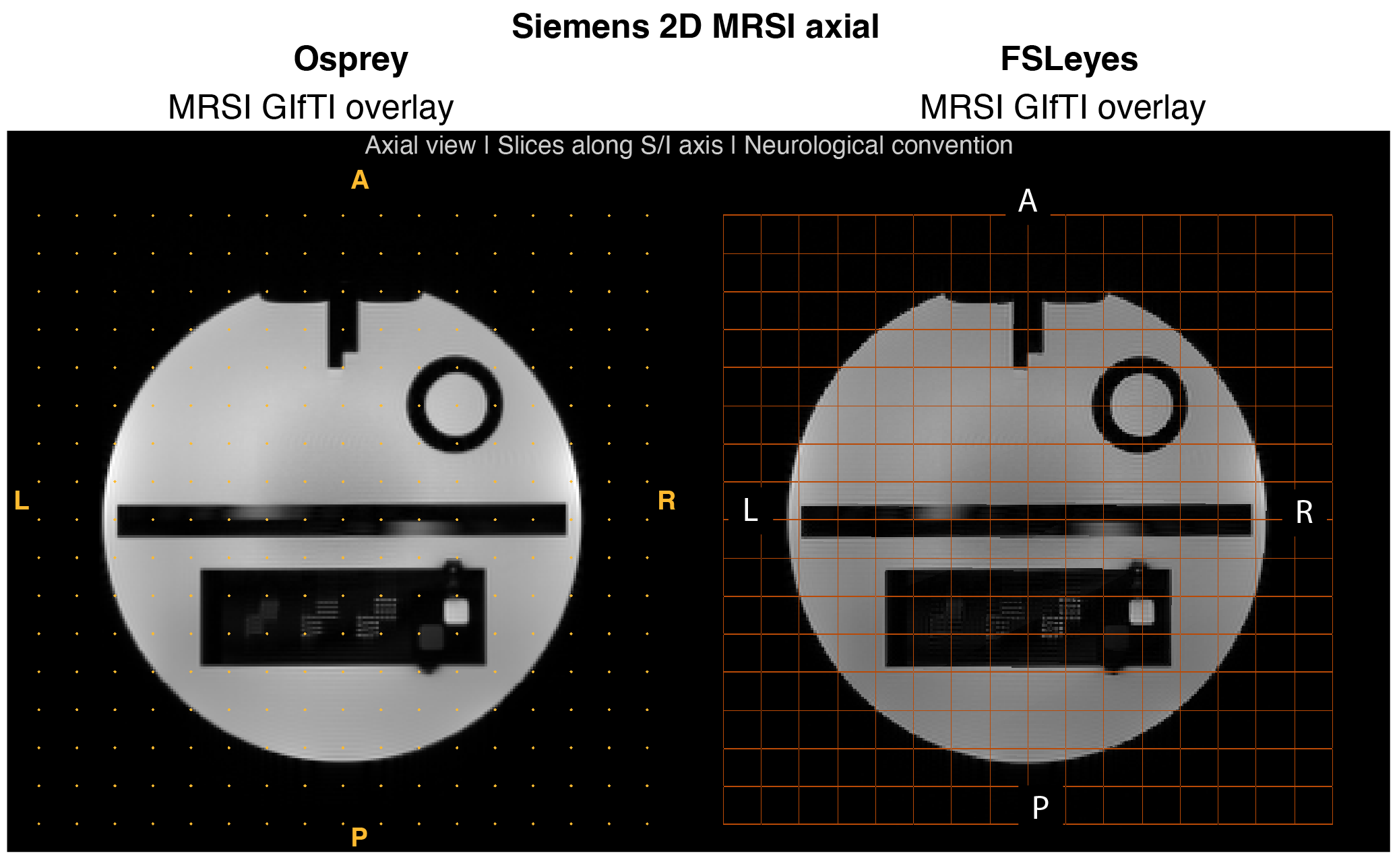

***Supplementary Material 7****. GE in vivo MRSI example datasets. Voxel localization and LCM results for the short- and medium-TE MRSI data from the workflow are shown.*

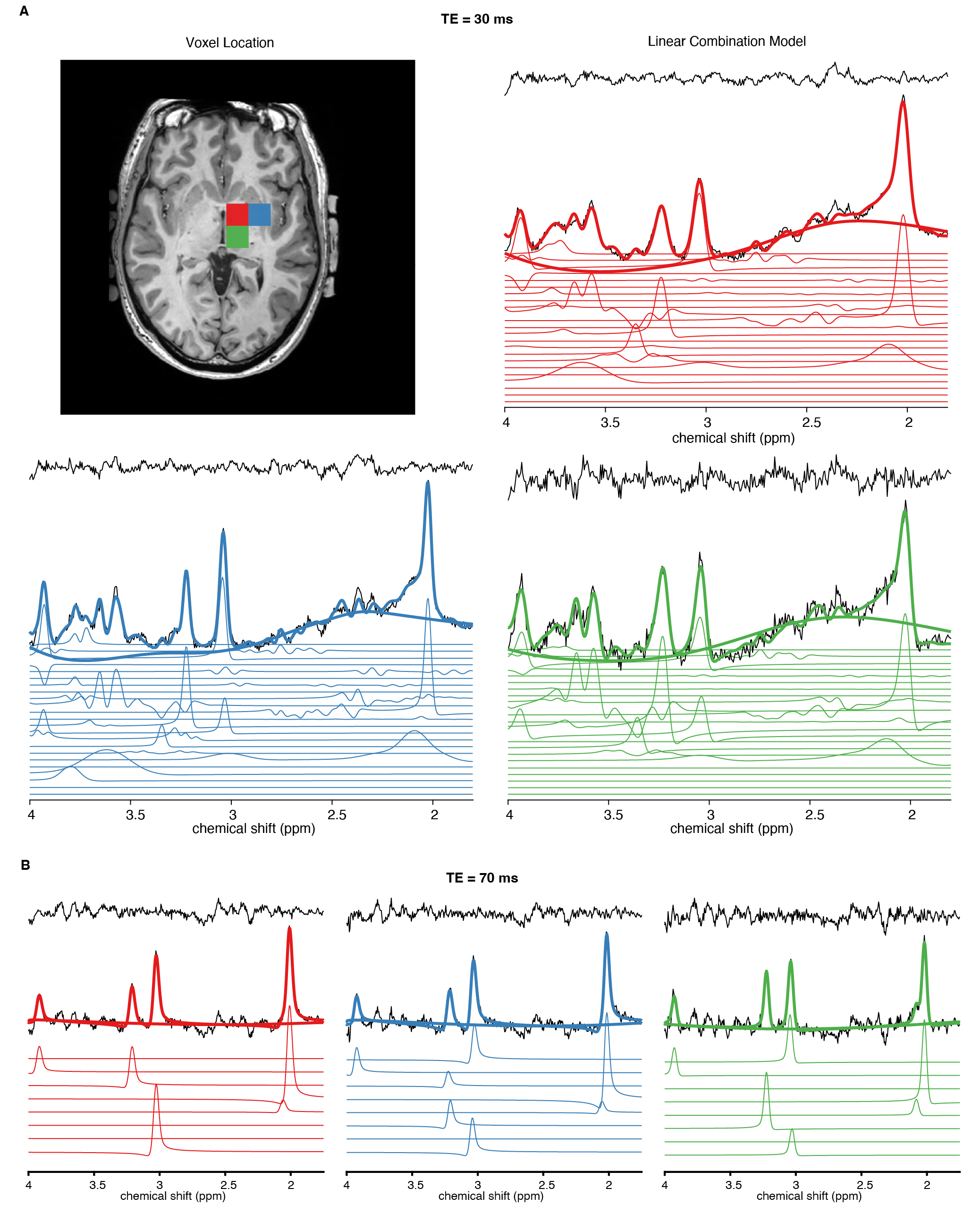

***Supplementary Material 8****. Siemens in vivo MRSI example datasets. Voxel localization and LCM results for the short- and long-TE MRSI data from the workflow are shown.*

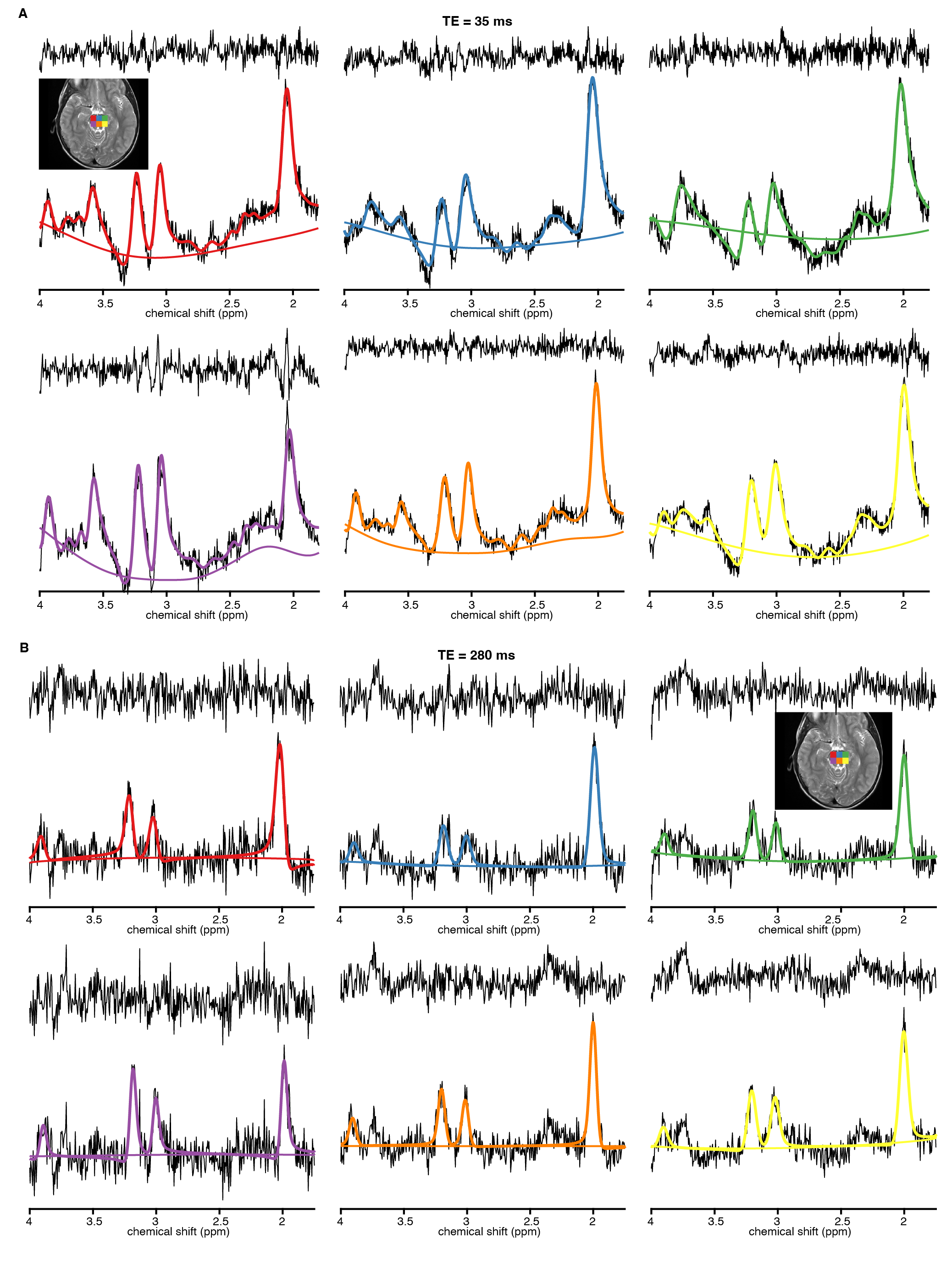

***Supplementary Material 9****. Example outputs from the semi-interactive HTML report. (A) Tissue fraction maps for gray matter (fGM), white matter (fWM), cerebrospinal fluid (fCSF), and lipids (fLip). (B) Quick map of the MRSI water reference scan generated by integrating the water signal. (C) MRSI example voxel locations and linear-combination model results from the corresponding locations. Note that the HTML report is originally semi-interactive, allowing the user to zoom into different parts of an image or spectrum or read the exact values (see the markers in each example). The HTML report contains, by default, visualizations of different quick maps, three spectra at each analysis stage from the center of the MRSI scan, and a visualization of QM. metrics and metabolites. Voxel locations and custom outputs can be modified by the user. An example of such a semi-interactive HTML file can be found on the GitHub repository.*

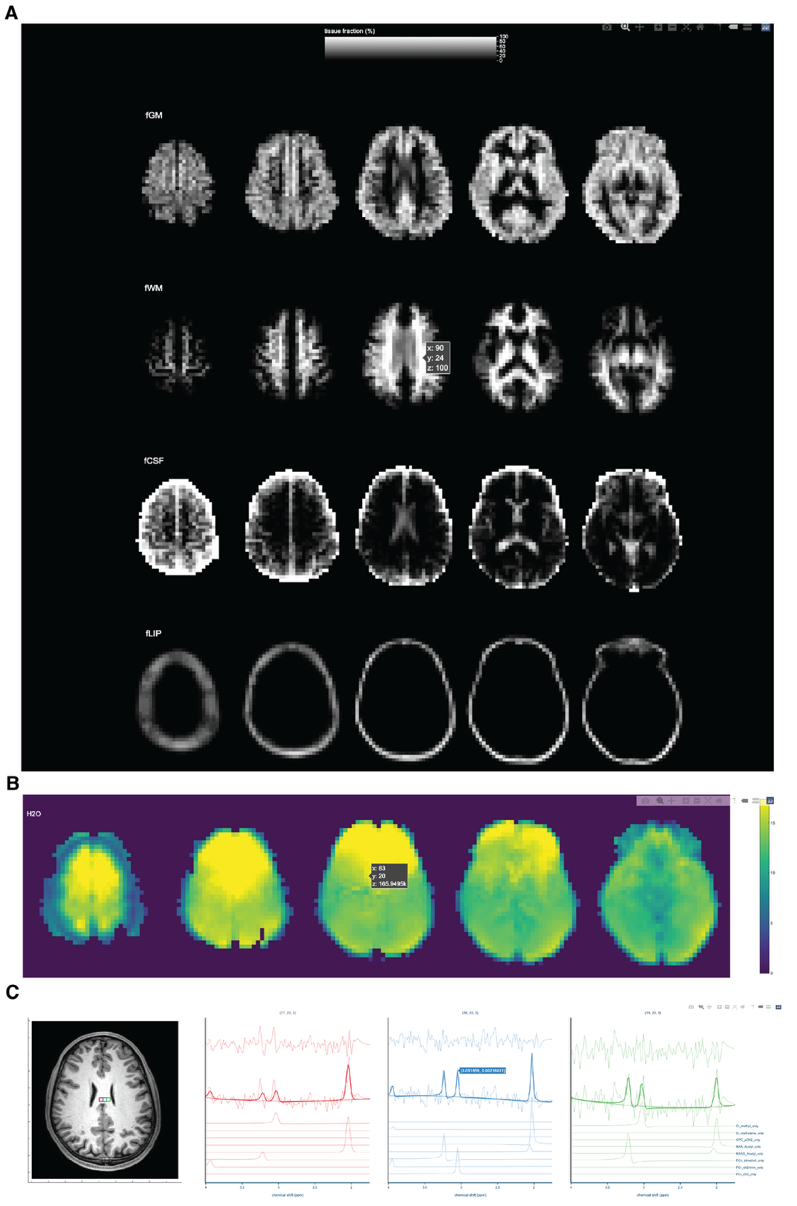
