## Supplementary material for "Fully automated open-source analysis and interactive visualization of magnetic resonance spectroscopic imaging (MRSI) data in Osprey-MRSI": Data Availability Statement

**Data and code availability statements**

The Osprey MRSI workflow is available in the Osprey-MRSI GitHub repository (<https://github.com/HJZollner/osprey-mrsi>). The scripts to run the analysis described in this manuscript are available in the example data folder (*/osprey-mrsi/exampledata)*. Run the workflow demonstration using <https://github.com/HJZollner/osprey-mrsi/blob/develop/exampledata/RunAllDatasets.m> and <https://github.com/HJZollner/osprey-mrsi/blob/develop/exampledata/ManuscriptFigures.m> to generate the raw figures. The Philips and Siemens datasets are available in the */osprey-mrsi/ exampledata/mrsi* subfolders. For GE, only the job files were shared on GitHub due to file size constraints. The user can download all example datasets, including the GE datasets, from the Center for Open Science Repository at <https://osf.io/vkdpr> and replace them in the */osprey-mrsi/ exampledata/mrsi* folder.
